## Supplementary figures and tables for "Binding to nucleosome poises human SIRT6 for histone H3 deacetylation"

### Supplemental Figures and legends

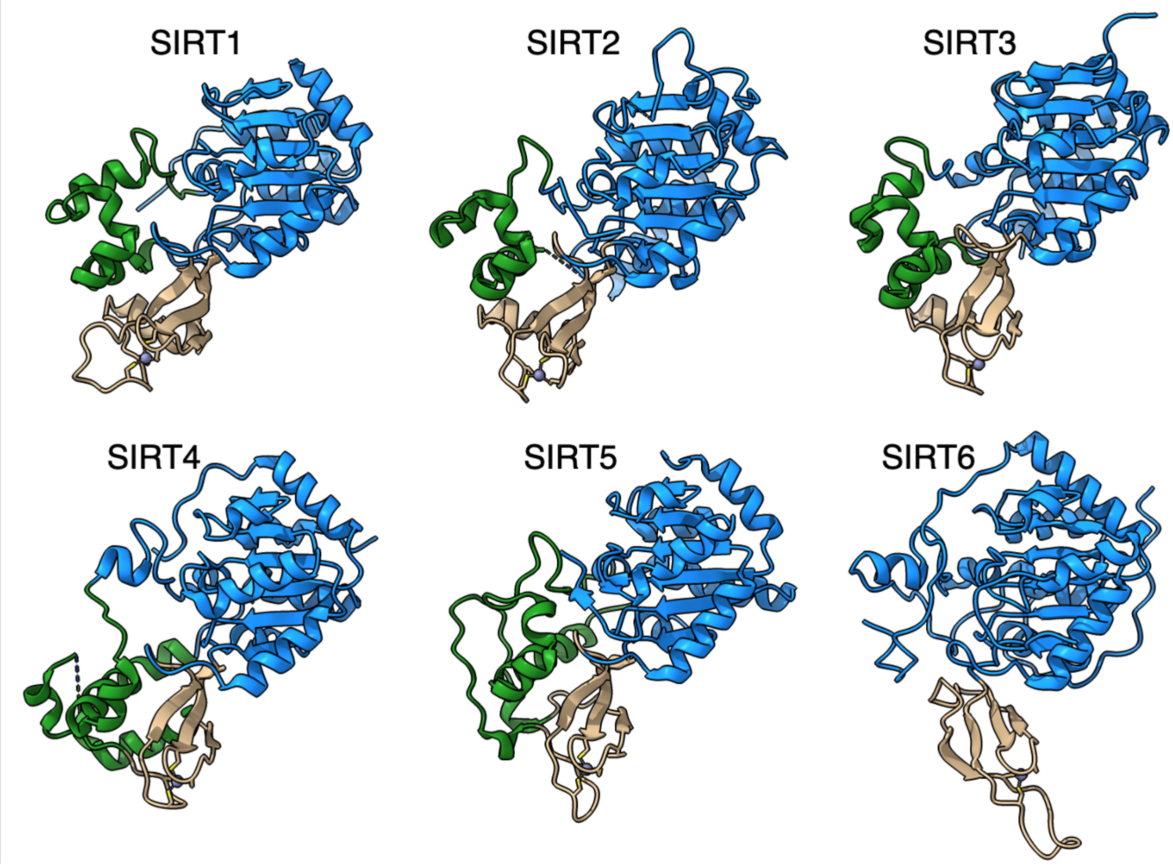


**Supplemental Figure 1 | SIRT6 lacks the helix bundle between the Rossmann fold and the Zinc-finger domains**

The structures of human sirtuins are depicted – the Rossmann fold domain in blue, Zinc-finger in tan and the helix bundle between the two in green.

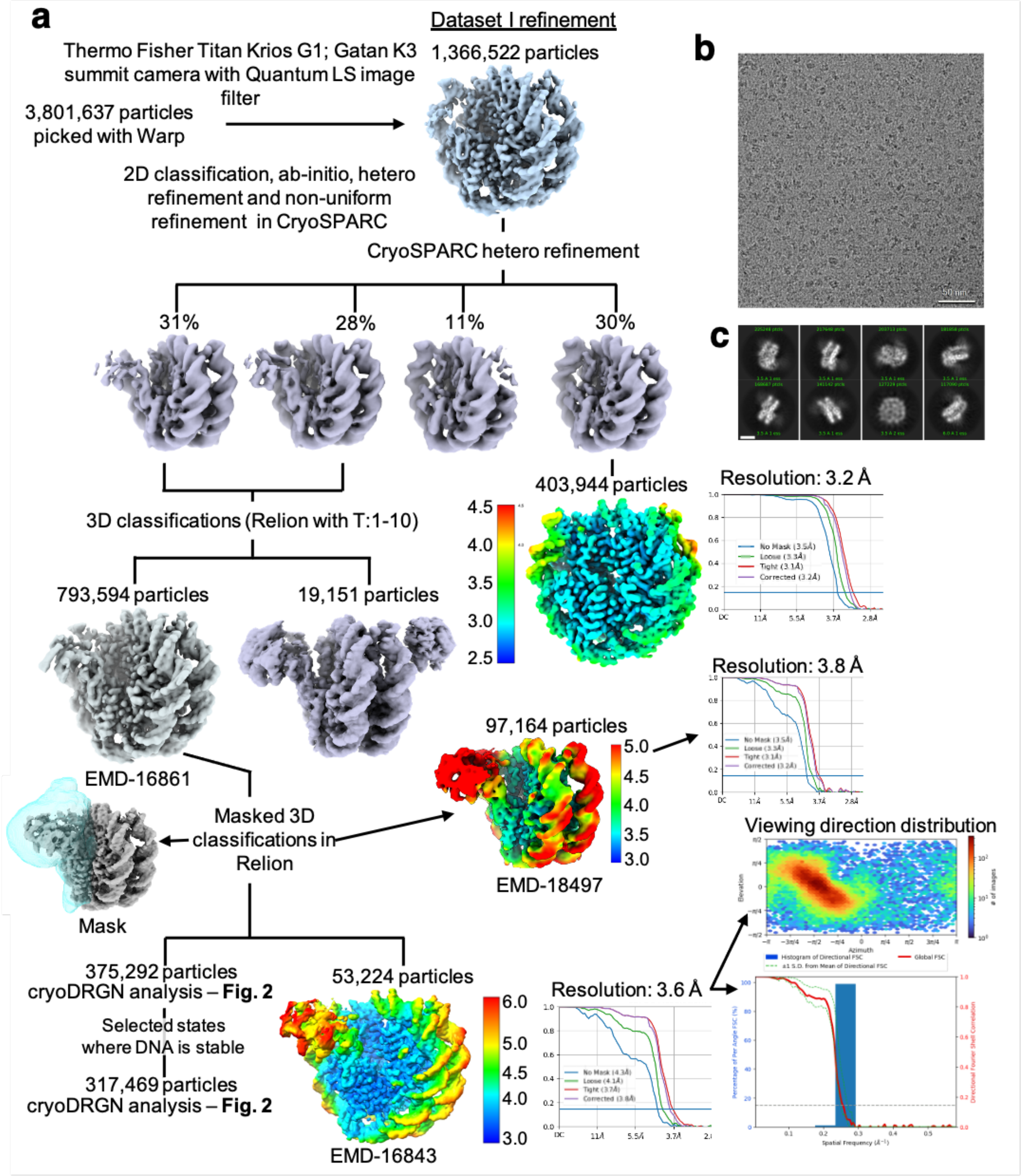


**Supplemental Figure 2 | Cryo-EM data analysis strategy for SIRT6-nucleosome – dataset 1**

**a**, Complete data processing scheme. Maps coloured in rainbow represent the local resolution of the reconstructions. Hollow blue volume represents the mask used for focused classification and refinement. FSC curves are depicted as a function of resolution in angstrom. CryoSPARC v.3 and v.4 were used to generate the 3.6 Å overall resolution map of SIRT6-nucleosome complex. RELION 3 was used for classifications of flexible regions, corresponding to the Rossmann fold domain of SIRT6. **b**, Original micrograph of SIRT6-nucleosome complex. **c**, Two-dimensional class averages showing high-resolution structural features.

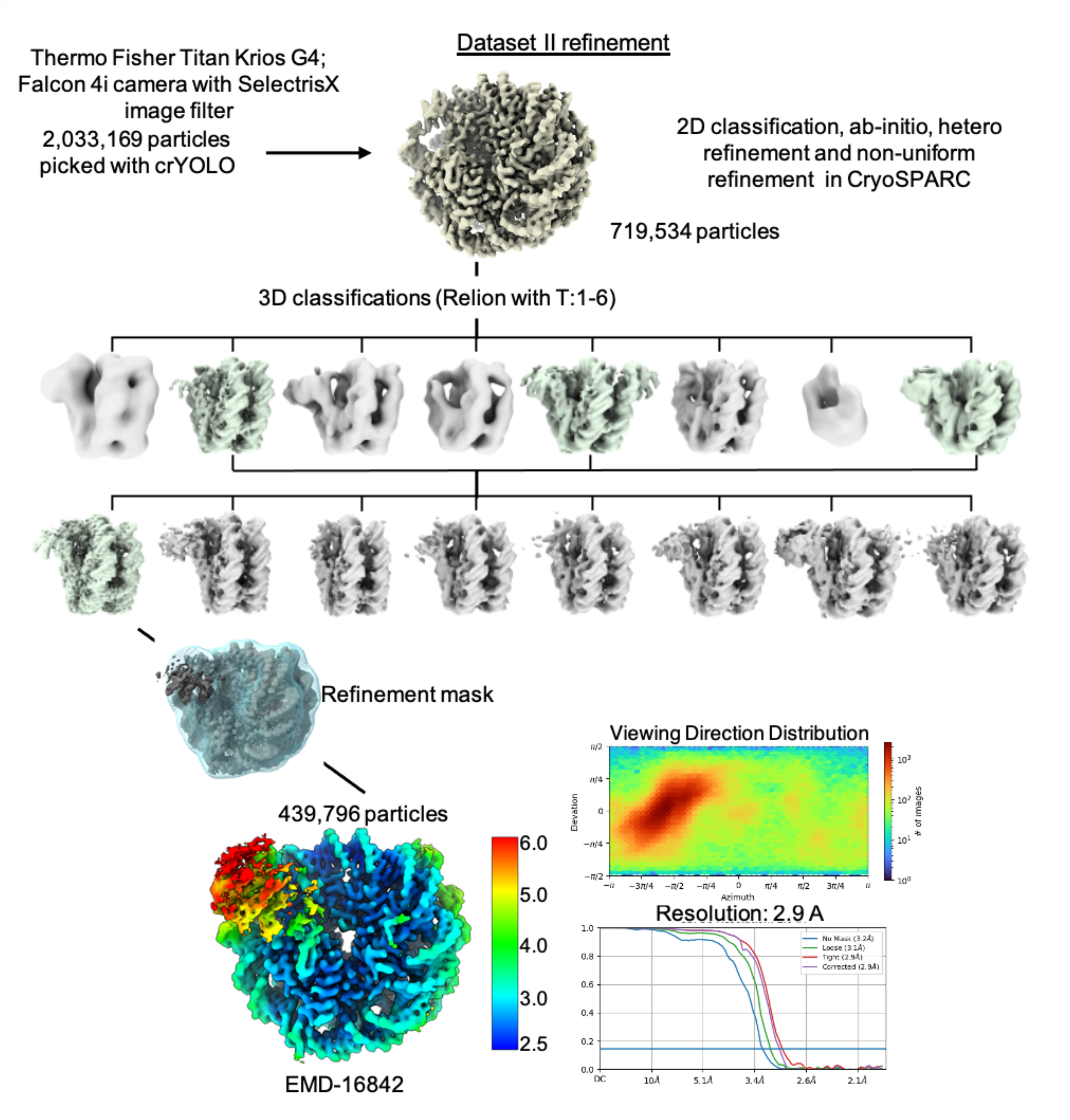


**Supplemental Figure 3 | Cryo-EM data analysis strategy for SIRT6-nucleosome – dataset 2**

**a**, Complete data processing scheme. Maps coloured in rainbow represent the local resolution of the reconstructions. FSC curves are depicted as a function of resolution in angstrom. CryoSPARC v.3 and v.4 and Relion 3 were used for 3D classification and to generate the 2.9 Å overall resolution map of SIRT6-nucleosome complex.

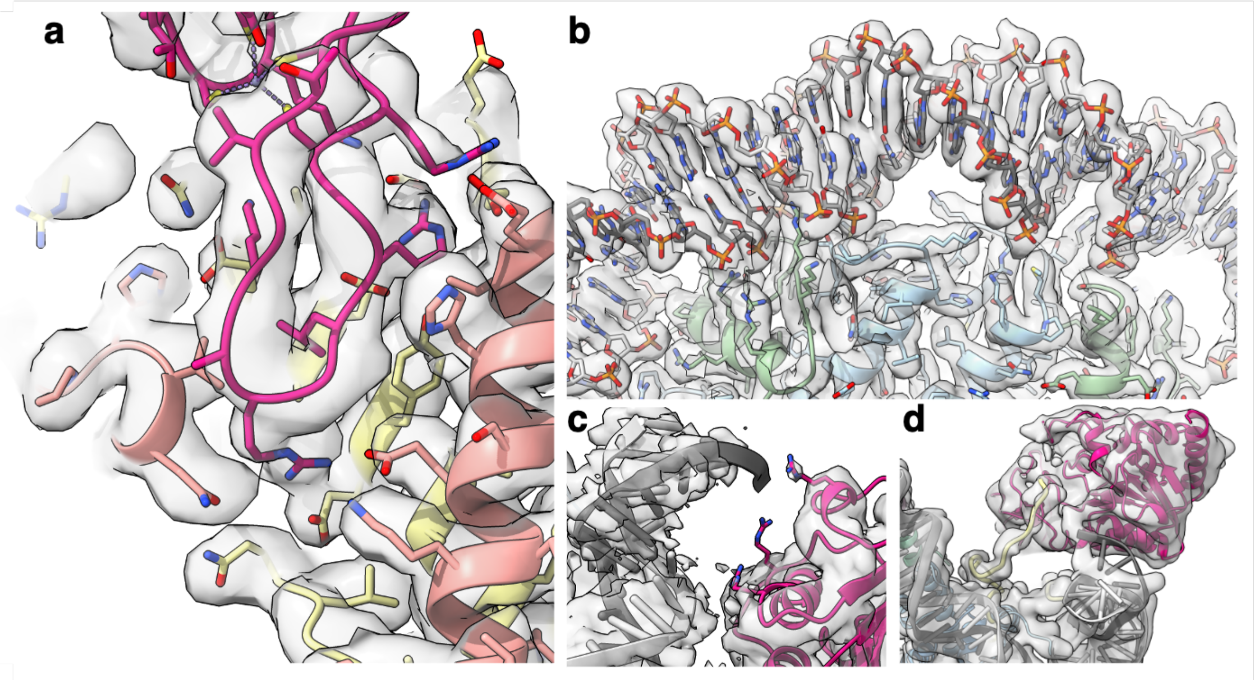


**Supplemental Figure 4 | Representative regions illustrating the quality of the cryo-EM map of SIRT6 bound to nucleosome**

**a**, Close-up view of the SIRT6 Zn-finger interacting with the nucleosomal acidic patch. **b**, View showing map details around the nucleosome dyad. **c**, SIRT6 Rossmann fold domain interaction with the DNA. **d**, Density used to trace H2A tail.

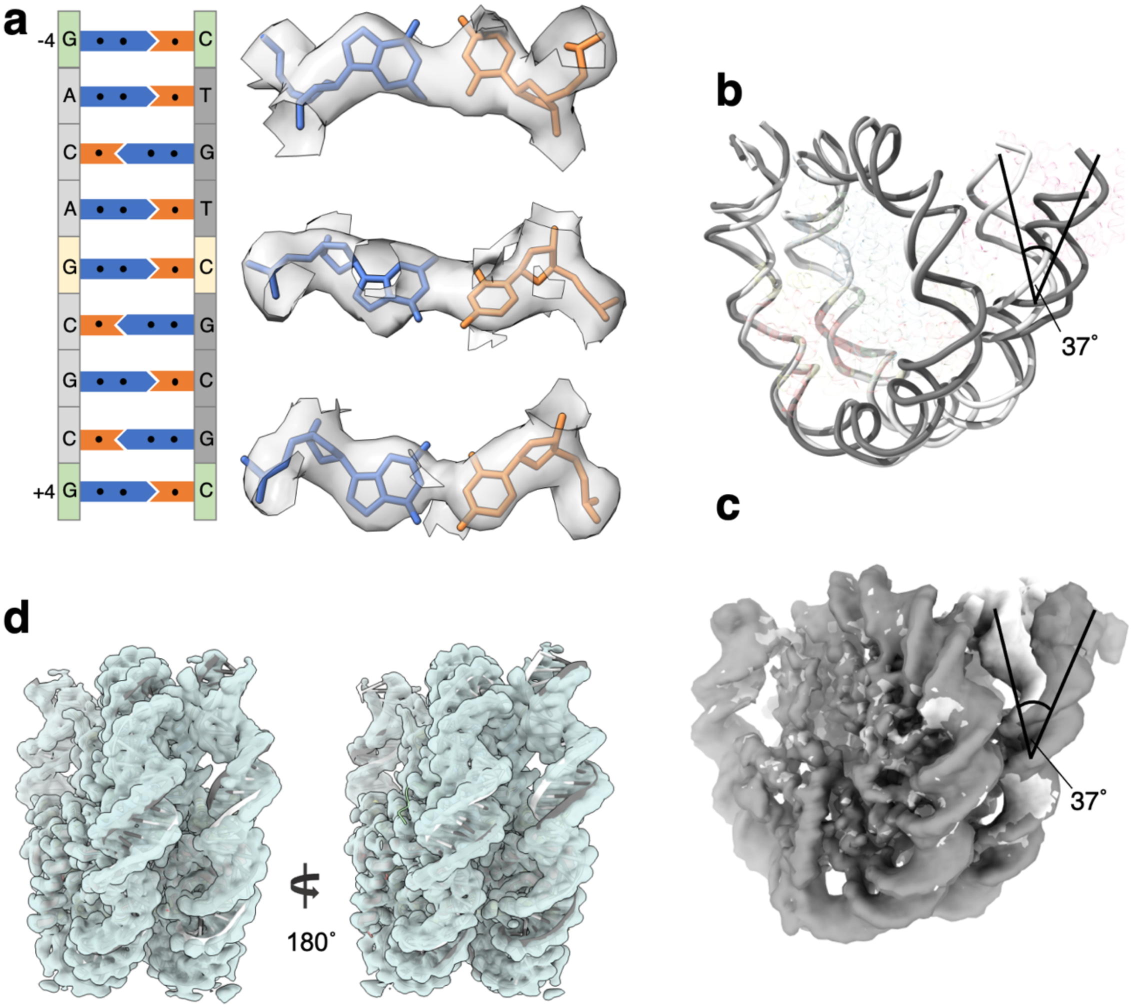


**Supplemental Figure 5 | SIRT6 binds to and displaces the "looser" DNA terminus**
**a**, Cryo-EM densities of the nucleotide base pairs at the dyad (yellow) and positions ± 4 (green) showing the orientation of the DNA in the SIRT6-nucleosome complex. Besides the dyad, the positions ± 4 are also asymmetric in the nucleosome with the Widom-601 DNA. **b**, The binding of SIRT6 to the nucleosome alters the path of the terminus DNA by 37°. Nucleosomal DNA is compared with (dim grey) and without (white) bound SIRT6. **c**, The same views as in panel b with the cryo-EM maps. **d**, Nucleosome without SIRT6 has no deviation of the DNA termini. PDB 3LZ0 was fitted to the nucleosome structure.

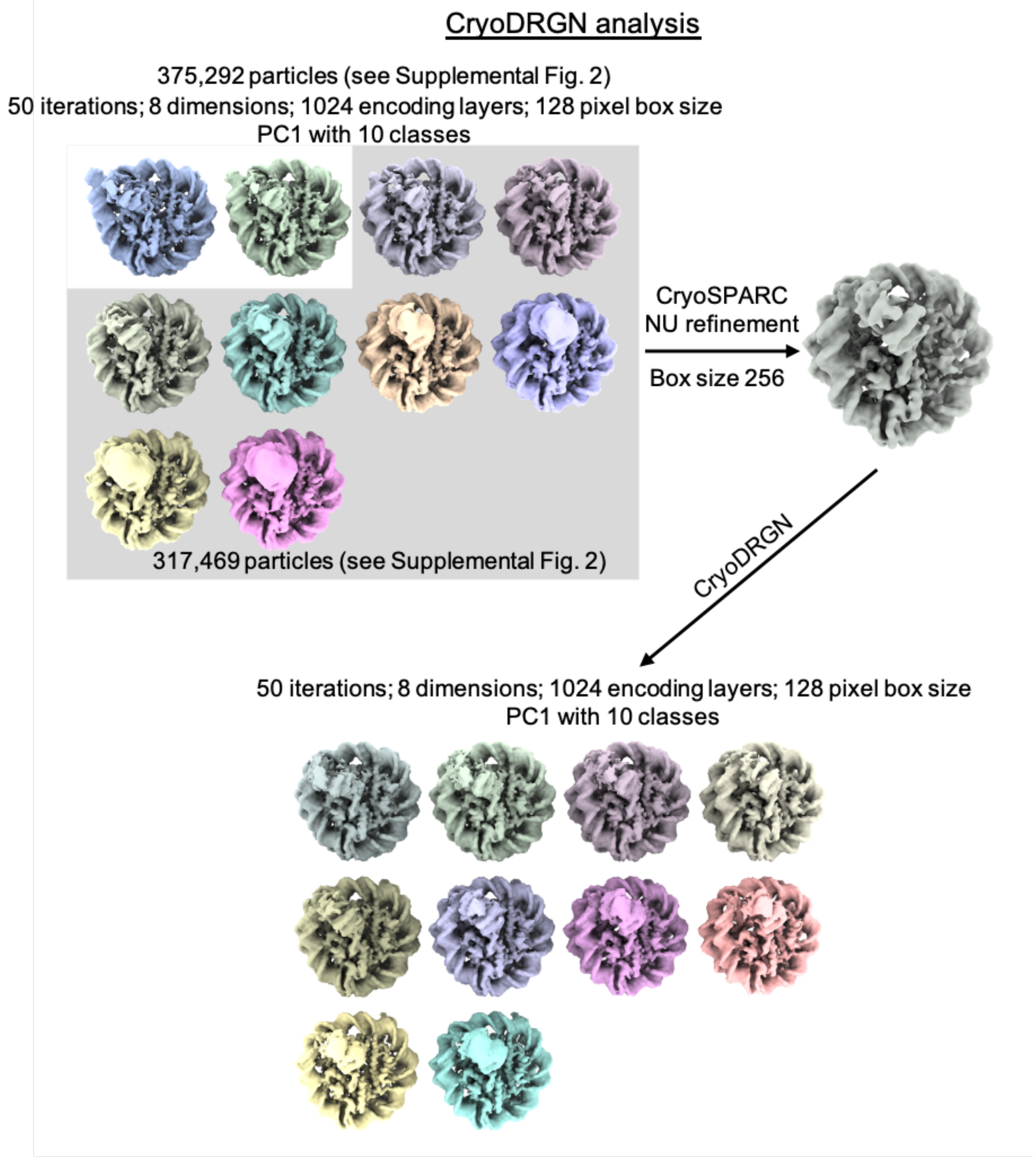


**Supplemental Figure 6 | Schematic representation of CryoDRGN analyses**
Particle images from CryoSPARC refinement were Fourier cropped to a size of 128x128 pixels. They were subjected to CryoDRGN analysis (run1). Ten representative volumes of the first Principal component is depicted. Particles from classes with similar DNA end positions were pooled and refined in CryoSPARC. Particles from this new refinement were re-analysed with CryoDRGN (run2). Ten representative volumes of the first Principal component is depicted.

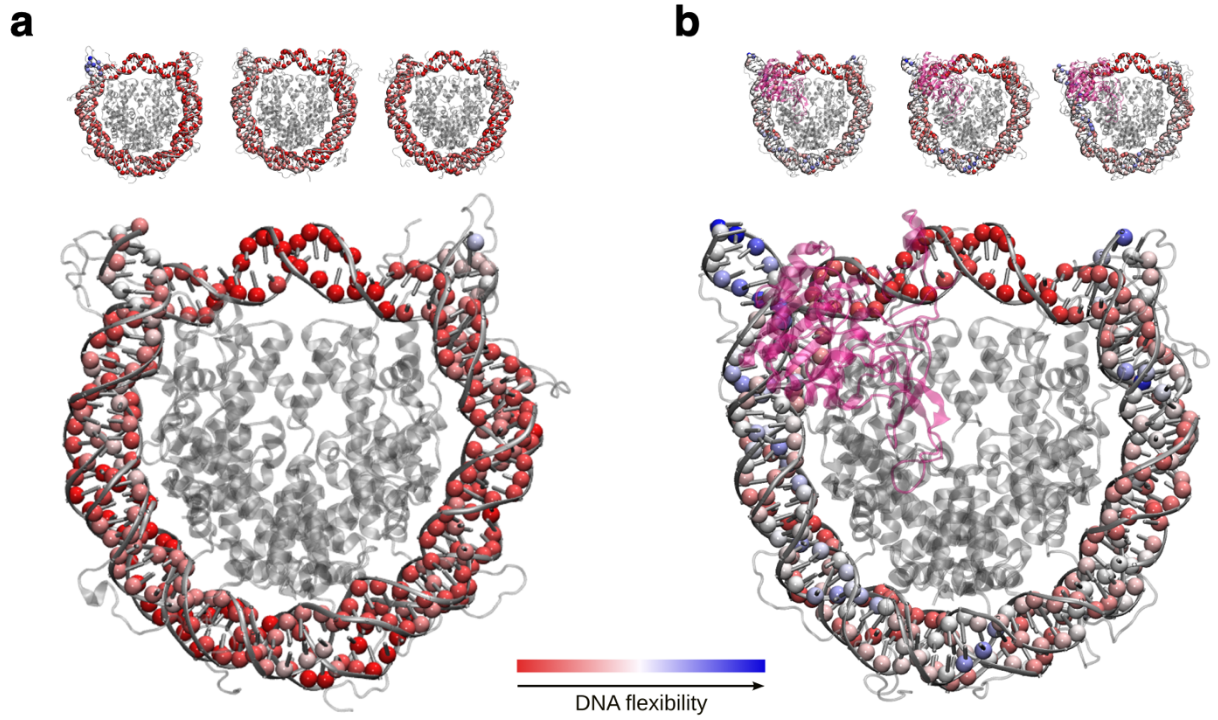


**Supplemental Figure 7 | SIRT6 binding renders DNA termini more flexible**
**a**, Projection of per residue flexibility contribution calculated from molecular dynamics simulations of the nucleosome without SIRT6. The balls at the DNA residues are colored according to the residue's relative contribution to the DNA overall flexibility. The 3 replicates are shown on the top and the projected average on the bottom. **b**, The same analysis is shown for the SIRT6 bound nucleosome. Rendered with VMD1.9.3.

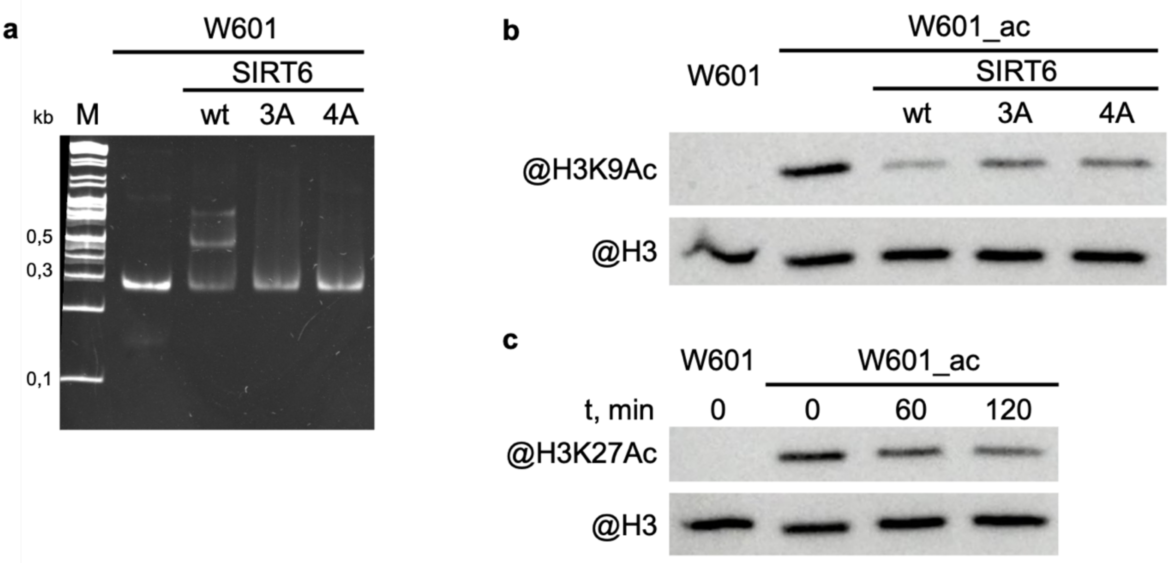


**Supplemental Figure 8 | WT and mutant SIRT6 - interactions with nucleosome and deacetylation activity**

**a**, Electrophoretic mobility shift assay to test binding of Widom-601 nucleosome to wt and mutant SIRT6. Lower and upper shifted bands visible in SIRT6 wt lane likely correspond to one or two SIRT6 molecules bound to one nucleosome. **b-c**, Deacetylation activity of SIRT6 and its mutants. W601_ac denotes SAGA-acetylated Widom-601 nucleosome. **b**, Western blot analysis of H3K9Ac deactylation by wt, 3A or 4A SIRT6. **c**, Western blot analysis of H3K27Ac deacetylation by SIRT6.

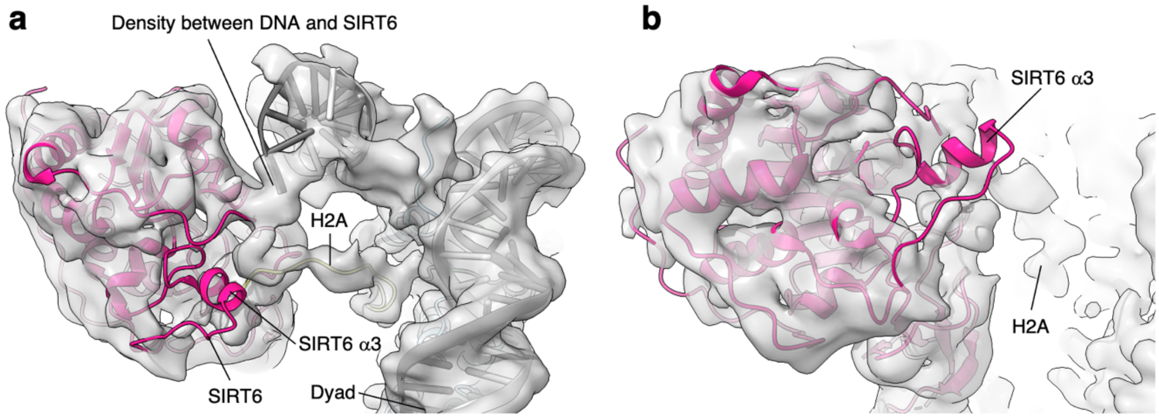


**Supplemental Figure 9 | SIRT6 rearrangement upon nucleosome binding**
**a**, Top view of the SIRT6 bound nucleosome. Two distinct densities were observed between SIRT6 and the DNA. **b**, Side view of the SIRT6 bound nucleosome. The third helix of SIRT6 (α3), which is part of the cofactor binding loop is not present in the structure.

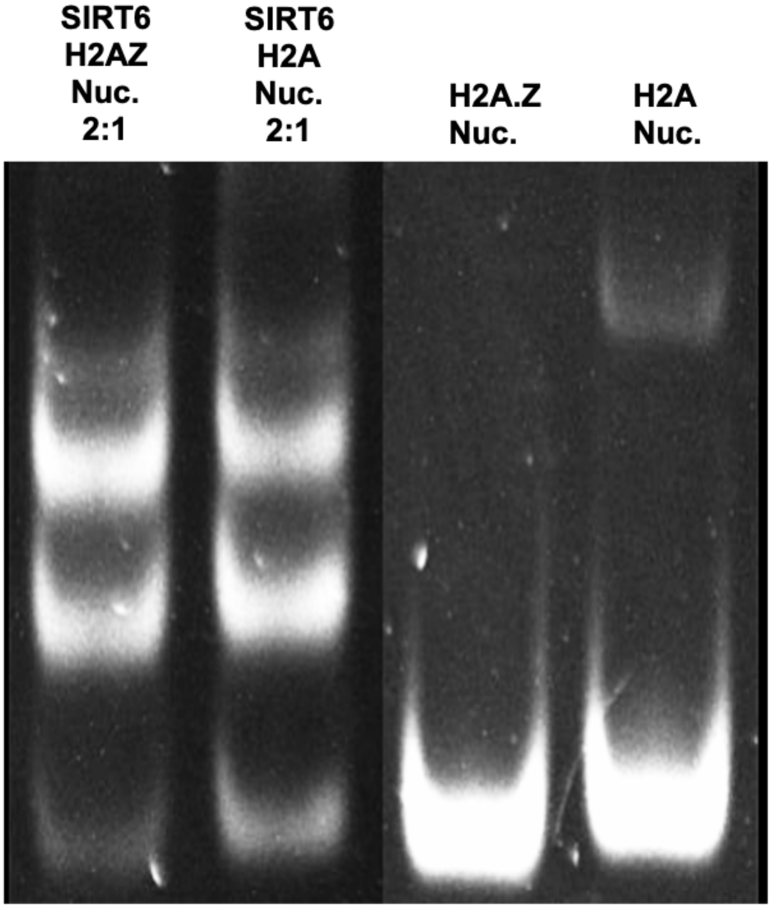


**Supplemental Figure 10 | Electrophoretic mobility shift assay comparing SIRT6 binding to H2A.Z containing nucleosomes and canonical (H2A containing) ones.**
Used for creating image 3f.

| | SIRT6-nucleosome #1 (EMDB-16843, EMDB-18497) | SIRT6-nucleosome #2 (EMDB-16842) | Composite map (EMDB-16845) (PDB 8OF4) |
|---|---|---|---|
| **Data collection and processing** | | | SIRT6 Rossmann fold: Dataset #1 Nucleosome and SIRT6 Zn-finger: Dataset #2 |
| Magnification | 81,000 | 270,000 | |
| Voltage (kV) | 300 | 300 | |
| Electron exposure (e–/Å$^2$) | 52 | 54.5 | |
| Defocus range (μm) | 1.2-3.4 | 0.8-2.6 | |
| Pixel size (Å) | 0.862 | 0.458 | |
| Symmetry imposed | C1 | C1 | |
| Initial particle images (no.) | 3,801,637 | 2,033,169 | |
| Final particle images (no.) | 53,224 | 439,796 | |
| Map resolution (Å) | 3.6 | 2.94 | |
| FSC threshold | 0.143 | 0.143 | |
| Map resolution range (Å) | 3.2-6.0 | 2.5-8.0 | |
| **Refinement** | | | |
| Initial model used (PDB code) | | | 3LZ0, 5X16 |
| Model composition | | | |
| Non-hydrogen atoms | | | 14,094 |
| Protein residues | | | 1,029 |
| DNA residues | | | 290 |
| Ligands | | | Zn: 1 |
| R.m.s. deviations | | | |
| Bond lengths (Å) | | | 0.013 |
| Bond angles (°) | | | 2.040 |
| Validation | | | |
| MolProbity score | | | 0.79 |
| Clashscore | | | 0.13 |
| Poor rotamers (%) | | | 0.58 |
| EMRinger score | | | 3.16 |
| Q-Score | | | 0.446 |
| Ramachandran plot | | | |
| Favored (%) | | | 96.43 |
| Allowed (%) | | | 3.57 |
| Disallowed (%) | | | 0.0 |

**Supplemental Table 1 | Cryo-EM data collection, refinement and validation statistics.**

| Cluster number | MD1 (5µs) | MD2 (5µs) | MD3 (5µs) |
|---|---|---|---|
| **With SIRT6** | | | |
| 1 | 32 % | 56 % | 37 % * |
| 2 | 23 % | 11 % | 11 % |
| 3 | 16 % | 6 % * | 10 % * |
| 4 | 7 % | 5 % * | 10 % * |
| 5 | 6 % * | 5 % * | 9 % * |
| 6 | 5 % | 5 % * | 7 % |
| 7 | 5 % * | 4 % * | 7 % |
| 8 | 3 % | 4 % * | 5 % * |
| 9 | 2 % | 3 % * | 3 % |
| 10 | 1 % | 1 % * | 1 % |
| **Without SIRT6 (control)** | | | |
| 1 | 34 % | 37 % | 34 % |
| 2 | 22 % | 17 % | 22 % |
| 3 | 9 % | 15 % | 16 % |
| 4 | 8 % | 11 % | 9 % |
| 5 | 8 % | 6 % | 8 % |
| 6 | 8 % | 4 % | 5 % |
| 7 | 5 % | 3 % | 4 % |
| 8 | 3 % | 3 % | 2 % |
| 9 | 2 % | 2 % | <1 % |
| 10 | 1 % | 2 % | <1 % |

**Supplemental Table 2 | Frequency of conformational clusters for each MD ensemble.** Clusters featuring the H3 tail protruding between the DNA and the octamer are marked by a star.
